# The motor pattern of rolling escape locomotion in *Drosophila* larvae

**DOI:** 10.1101/2022.11.03.514605

**Authors:** Liping He, Lydia Borjon, W. Daniel Tracey

## Abstract

When undisturbed, *Drosophila* larvae move forward through their environment with sweeping waves of caudal to rostral muscle contraction [1, 2]. In stark contrast, nociceptive sensory stimuli (such as attacks by parasitoid wasps) trigger the larvae to roll across the substrate by corkscrewing around the long body axis [3, 4]. While studies have described the motor pattern of larval crawling [1, 2], the motor pattern of larval rolling escape locomotion remains unknown. Here, we have determined this pattern. To do so, we developed a high speed confocal time-lapse imaging preparation that allowed us to trigger rolling with optogenetics while simultaneously imaging a genetically encoded calcium sensor that was expressed in the muscles. Of the 30 muscles present in each larval abdominal hemisegment we find that only 11 muscles are consistently and specifically activated across segments during rolling. 8 additional muscles are more sparsely activated. Importantly, the sequential pattern of muscle recruitment during rolling is completely distinct from that of forward or reverse crawling. We discover that a roll involves a wave of muscle activation that propagates around the larval circumference (in the transverse plane of each segment) and involves four coactive muscle groups. A pattern of activation progresses from coactive ventral muscle groups to dorsal groups and then spreads across the midline to the contralateral dorsal muscle groups which then progresses back to the ventral groups. Finally, the direction of a roll (either clockwise or counterclockwise around the body) is determined by the clockwise or counterclockwise order of muscle group activation around the transverse plane.

## Results and Discussion

In order to observe fluorescent calcium signals indicative of muscle activation we imaged rolling second instar larva through a coverslip on an inverted confocal microscope (**Figure 1A**). An agar pad with a water filled cutout depression atop the coverslip was used to restrict the movement of the larva to the field of view. The blue 488 nm laser allowed for excitation of GCaMP6.0f [5] under the control of a muscle specific driver (R44H10-LexA) [2]. The blue laser simultaneously triggered rolling through optogenetic activation of nociceptive class IV multidendritic arborization neurons since our experimental larvae expressed Channelrhodopsin2∷YFP in those cells (under control of *pickpocket-GAL4*) [4]. We generated time-lapse recordings under these experimental conditions and tested larvae were seen to roll in a random direction (ie. either towards the right side of their body (clockwise) or towards the left side (counterclockwise)) **(Figure 1B)**. We obtained 19 recordings (9 clockwise and 10 counterclockwise) with complete rolls which were used for further analysis. The duration of a rolling cycle (ie. amount of time required for the larvae to complete a full roll) was 1.60±0.36 seconds (mean±SD, n=9) for clockwise rolling and 1.7±0.46 seconds (mean±SD, n=10) for counterclockwise rolling.

**Figure 1:**
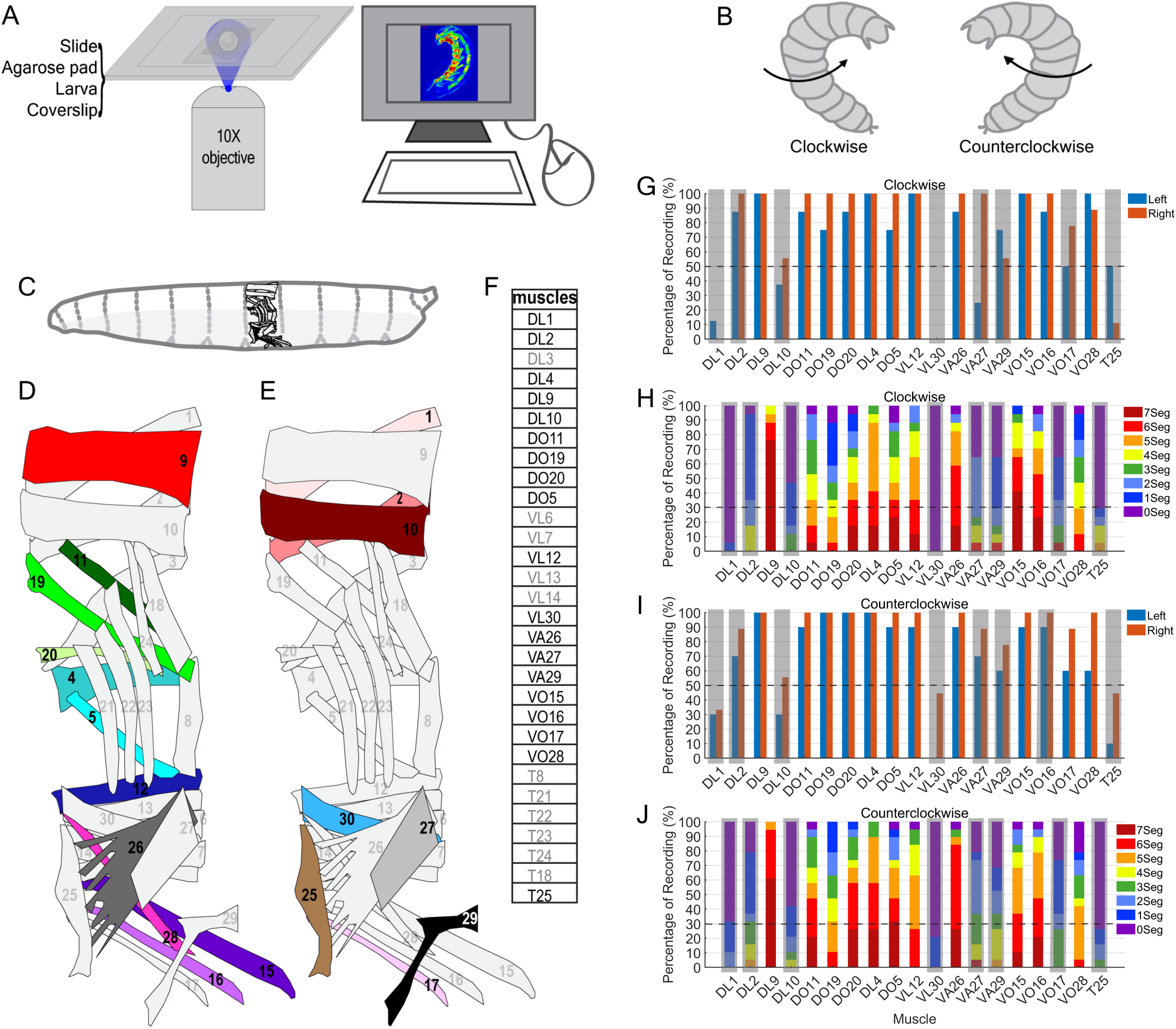
The abdominal muscles that are activated during larval rolling behavior. (A) Schematic of the imaging preparation. The larva was imaged on an inverted confocal microscope through a coverslip. An agar pad above the larva contained a water filled depression and was mounted on a glass slide. (B) Schematic drawing of clockwise and counterclockwise rolling, heads of the larvae are up, the tails are down towards the bottom of the image. (C) Schematic of a Drosophila larva as viewed from the side. 30 body wall muscles are shown in the 5th abdominal hemisegment. (D) Muscles that were most commonly activated during larval rolling. Solid colors indicate muscles that were consistently activated in our recordings. (E) Muscles that were sparsely or rarely activated during larval rolling. Solid colors indicate the activated muscles. (F) List of the 30 abdominal muscles with the activated muscles shown in bold font. (G) The frequency of recordings where activation was observed is graphed for each of the muscles involved in clockwise rolling. (H) Plot of the number of segments that showed activation for each muscle across recordings (ranged from 0 to 7) for clockwise rolling. (I) The frequency of recordings where activation was observed is graphed for each of the muscles involved in counterclockwise rolling. (J) Plot of the number of segments that showed activation for each muscle across recordings (ranged from 0 to 7) for counterclockwise rolling.

Each abdominal segment contains 30 muscles on each side of the body (30 on the left and 30 on the right). A well-trained investigator can identify each of these muscles according to their morphologies, their position on the dorsal ventral axis, and their position in relation to other nearby muscles. Using careful frame-by-frame analysis we painstakingly scored the activation of individual muscles in abdominal segments 1-7 (A1-A7) during rolling. If we observed that the GCaMP6.0f fluorescence signal in the muscle became obviously brighter during the roll we scored it as activated. This determination of muscle activation was a qualitative/binary ON or OFF designation as our imaging conditions did not allow for quantitative determination of changes in fluorescence intensity. However, the strong magnitude of the GCaMP6.0f fluorescence changes during our recordings made the ON determination clear and unambiguous (see Supplemental video 1, 2). A caveat of this methodology is that small quantitative changes in the calcium indicator were ignored, and thus we cannot eliminate the possibility that we have missed weakly activated muscles in our analysis.

### A subset of muscles is involved in rolling

From this analysis, we found that 11 of the 30 muscles in each abdominal hemisegment were consistently and robustly activated across recordings in both clockwise and counterclockwise rolling behaviors (**Figure 1 C-J**). These 11 muscles included dorsal longitudinal muscles (DL) 9 and 4, dorsal oblique muscles (DO) 11, 19, 20 and 5, the ventral longitudinal muscle (VL) 12, ventral acute muscles (VA)26 and ventral oblique muscles(VO) 15,16 and 28. We observed that these muscles were each activated in 3 or more body segments in over 70% of recordings. Two muscles (DL2 and VA29) were also activated in greater than 70% of recordings but in a sparser and segment specific pattern that was rarely seen in greater than 3 segments (**Figure 1G-J**). An additional 6 muscles were more rarely activated (DL1, DL10, VL30, VA27, and T25) (**Figure 1 D-J**). Note that we observed activation of muscles VA27 and VO17 in the majority of recordings but due to their close proximity to other strongly activated muscles we could not unambiguously score their activity. Interestingly, although we made a focused effort, we did not observe activation of the transverse muscles 21, 22 or 23 which have been previously proposed to be important to rolling behavior [6, 7] despite having verified that the R44H10-LexA driver is robustly expressed in those muscles (data not shown, [2]). However, we cannot exclude the possibility that these muscles are weakly activated during rolling, or that they are activated at a phase of the rolling cycle that was difficult to observe (ie. in the C-Bend, see below). It is interesting to note that 8 of the 11 commonly activated rolling muscles show an angled (acute or oblique) anatomy while only 3 of these muscles are longitudinal. This latter observation likely has important implications for the physical basis of rolling locomotion as shortening of these muscles is predicted to create tension in both the longitudinal and the dorsal ventral axes.

### A sequence of muscle contractions along the dorsal ventral axis occurs during clockwise rolling

We next determined the timing of the calcium increases for each of the muscles involved in rolling. In order to compare the recordings of different rolling events, we established anatomical landmarks from which to determine a nomarlized time scale (**Figure S1**). We defined the beginning of a rolling cycle (normalized time 0) to be the point during a roll when the ventral midline was precisely centered relative to the left right axis. The midpoint of a rolling cycle (normalized time 0.5) was defined as the point when the dorsal midline was precisely centered. The end of the cycle (normalized time 1) was when the ventral midline was again centered. Note that a roll actually initiates prior to the time 0 of the rolling cycle and the normalization is purely to allow comparison of different rolls which are of varying time durations.

We carefully observed frame-by-frame still images from our recordings and noted the timestamp for the frame of the recording when each muscle first became activated in abdominal segments one through seven. We also noted the duration of activation for each muscle during the period that it was clearly visible and identifiable in the recordings. It was not possible to keep track of individual muscles for the entire rolling cycle for two reasons. First, our imaging conditions only allowed us to view the larvae from a single angle and the depth of field under our imaging conditions allowed us to observe approximately half of a larva (in the sagittal/longitudinal plane) at any given time. Second, when the muscles contracted very strongly, this caused the larva to form the characteristic C shape (or C-bend) that is typical of rolling behavior [3, 8]. The strongly contracted muscles of the C-bend blended together and thus could no longer be resolved from one another which did not allow for unambiguous determination of where the calcium signals originated.

Nevertheless, we were able to resolve the sequence of muscle activation for both clockwise and counterclockwise rolling because muscles clearly and strongly brightened during activation. At the start of a rolling cycle a prominent ventral flexion (primarily involving muscles VO15R, VO16R, VO26R and VO28R) was observed (**Figure 2A-D**). During clockwise rolling this ventral flexion spread dorsally on the right side of the body with recruitment of muscles VL12R, DO5R, and DL4R and eventually spreading to DO20R, DO19R and finally to muscle DL9R (**Figure 2A,B**). The pattern of muscle contraction then crossed the dorsal midline to recruit muscles from the left side of the body beginning with the dorsal muscles and spreading ventrally in the opposite order to the pattern that happened on the right side of the body. The longitudinal DL9L was followed by DO11L, DO19L, DO20L, then DL4L, VL12L, VA26L and finally VO28L, VO15L and VO16L (**Figure 2A,B).**In Figure 2C we plot the detailed timing of activation of muscles seen across segments A1-A7 for the representative recording shown in Figure 2 A and B. In this plot it can be clearly seen how the pattern of contraction spreads from ventral to dorsal, crosses the midline, and then spreads from dorsal to ventral. To seek a characteristic average pattern of muscle recruitment across all of the clockwise rolling recordings we plotted the number of activated segments versus the normalized time axis of the rolling cycle (**Figure 2D**). Although the high-speed analysis of individual recordings allowed us to see individual muscles turning on in a sequence, the merged analysis across recordings suggests 4 coactive groups of muscles on each side of the body that are sequentially recruited during the rolling cycle. A ventral group of four muscles (VO15, VO16 VA26 and VO28), a ventral-lateral group of three muscles (VL12, DL4 and DO5), a dorsal-lateral group of three muscles (DO11, DO19, and DO20) and finally a dorsal group with one or 2 muscles (DL9 and/or DL2) (**Figure 2D)**. Note that although DL2 is activated in most of our recordings, it was sparsely activated in a subset of segments (Figure 1) and it is not plotted in Figure 2D for clarity.

**Figure 2:**
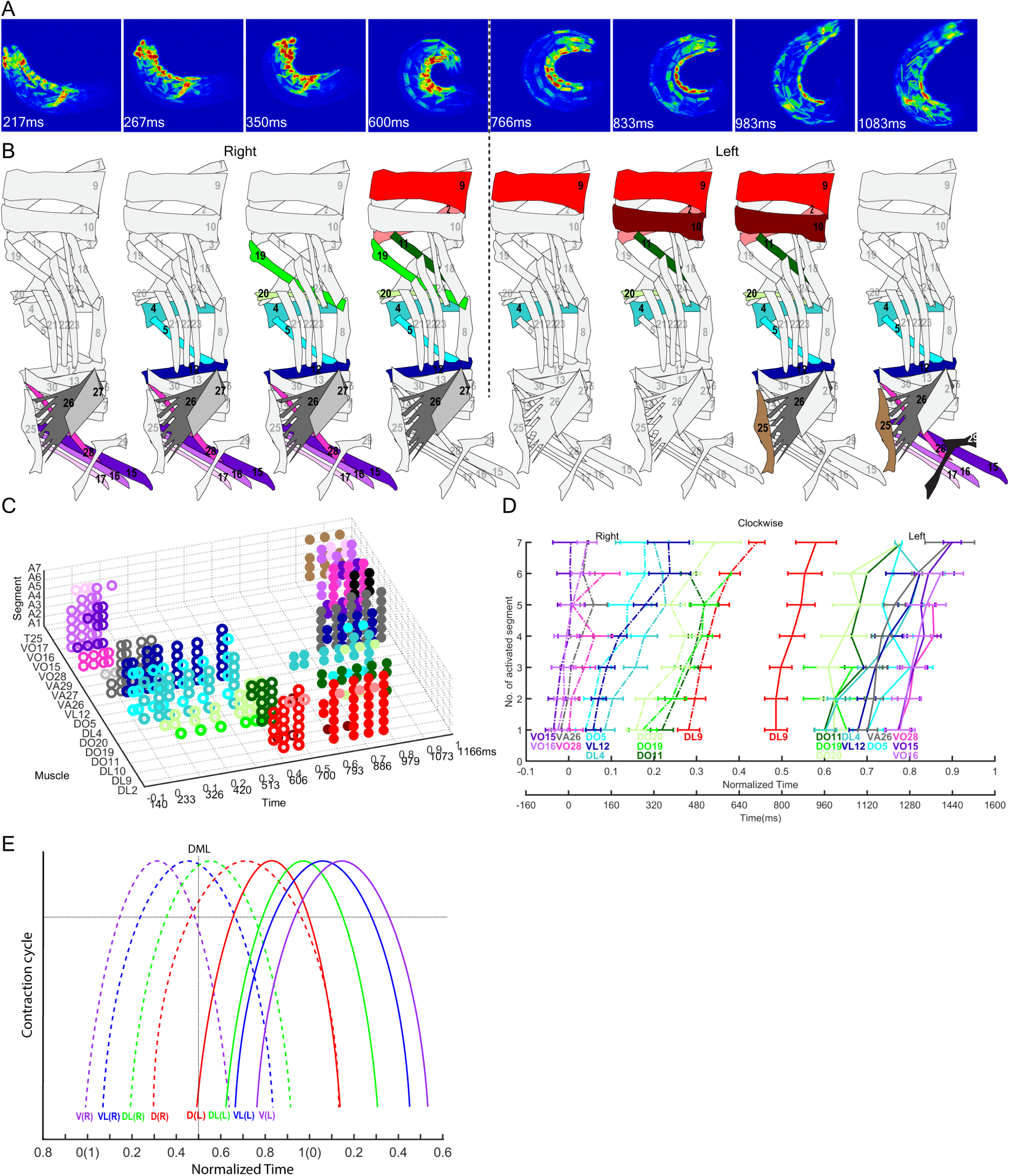
The sequence of muscle recruitment during clockwise rolling. (A) Selected frames from a recording of muscle activity during clockwise rolling behavior. Colors from blue to red represent the intensity of GCaMP6f signal from low to high (See Video S1). (B) Schematic representation of the visibly activated muscles in larval abdominal hemisegments for the frames shown in panel A. Solid colors indicate the activated muscles that are visible in the frame above. (C) Scatter 3D plot depicting the sequence of activated muscles across the abdominal segments during clockwise rolling. The Z axis shows the seven segments, the Y axis indicates the identity of individual muscles, the X axis shows the time normalized to the rolling cycle as well as the true time from the recording. (See Video S3 for the recording of this rolling event). (D) Plot of the average recruitment timing for the 11 abdominal body wall muscles that are activated during larval clockwise rolling. Error bars represent the standard errors (n=9). The X axis labels show the normalized rolling phase and the average Min-Max normalized time respectively. (E) Schematized estimate of the contraction cycle for each muscle group. The peak of each curve indicates the point in the contraction cycle when each muscle group is located within the strongly contracted C bend. The ventral (V) group includes VO15, VO16, VA26 and VO28. The ventral lateral (VL) group includes VL12, DO5 and DL4 muscles. The dorsal lateral (DL) group includes DO11, DO19 and DO20. The dorsal group includes only muscle DL9. (L) indicates the left hemisegment, (R) denotes the right hemisegment. The horizontal dashed line denotes the approximate point in the contraction cycle where GCaMP6f signals are detectable in all 7 abdominal segments for the muscle group.

The point in Figure 2D where a given muscle is activated in all seven segments represents the point in the rolling cycle where calcium signals have clearly risen above baseline across segments. These signals also reflect the point where all seven segments have initiated contraction but they do not inform directly on the magnitude of muscle shortening. We observed that muscle shortening obviously continued for all of the muscles into the C-bend, but the absolute magnitude of the contraction could not be measured since the muscles could not be resolved within the C-bend. However, using the muscle recruitment data from Figure 2C we were able to estimate the pattern of contraction and relaxation for each of the 8 coactivated muscle groups (**Figure 2E**).

### The sequence of muscle contractions in counterclockwise rolling is the reverse of clockwise rolling

During counterclockwise rolling, muscle contractions initiate on the opposite side of the body relative to clockwise rolling. Thus, the ventral flexion of counterclockwise rolling propagates first to the left side of the body which is the opposite of what was seen in clockwise rolling (**Figure 3A-D**). The rolling sequence is initiated with ventral muscles VO15L, VO16L, VA26L and VO28L (**Figure 3A,B)**. This is followed by VL12L, DL4L and DO5L then DO11L, DO19L, and DO20L and eventually DL9L. As with clockwise rolling, counterclockwise rolling spreads across the dorsal midline but engages DL9R followed by dorsolateral (DO20R, D19R, DO11R), ventrolateral (DO5R, DL4R, VL12R) and eventually the ventral rolling group (VO15R, VO16R, VA26R, VO28R) (**Figure 3A,B)**. In Figure 3C, we plot the detailed timing of activation of muscles seen across segments A1-A7 for the representative recording shown in Figure 3 A and B versus time. Again, it is clearly evident that the pattern of activation progresses from ventral to dorsal, spreading across the midline, and then progressing from dorsal to ventral. Analysis of the averaged pattern of activation across recordings suggests that the same four coactive muscle groups that were activated during clockwise rolling were apparent with counterclockwise rolling (**Figure 3D**). But the order of activation for these groups was the opposite of what was seen with clockwise rolling (**Figure 2D, Figure 3D**). Similarly, our estimated pattern for contraction and relaxation cycle for these muscle groups appears to be the mirror image of that for clockwise rolling (**Figure 3E**).

**Figure 3:**
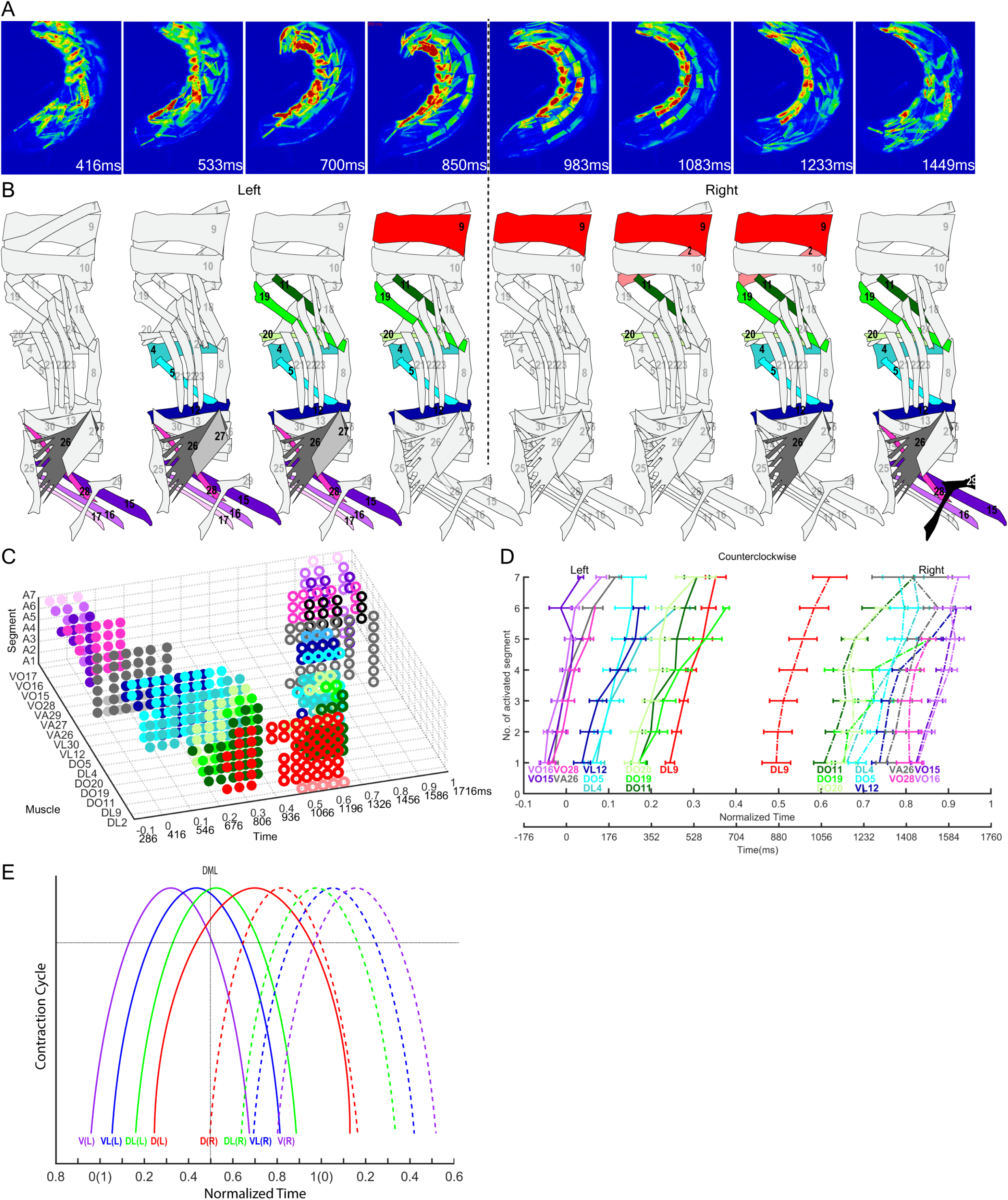
The sequence of muscle contraction during counterclockwise rolling. (A) Selected frames from a recording of muscle activity during counterclockwise rolling behavior. Colors from blue to red represent the intensity of GCaMP6f signal from low to high (See Video S2). (B) Schematic representation of the visibly activated muscles in larval abdominal hemisegments for the frames shown in panel A. Solid colors indicate the activated muscles that are visible in the frame above. (C) Scatter 3D plot of activated muscles in larval abdominal segments during a ventral-to-ventral counterclockwise rolling. The first and second X axis are the normalized rolling phase and the real time respectively (See also Video S4). (D) Plot of the average recruitment timing for the 11 abdominal body wall muscles that are activated during larval counterclockwise rolling. Error bars represent the standard errors (n=10). The X axis labels show the normalized rolling phase and the average Min-Max normalized time respectively. E) Schematized estimate of the contraction cycle for each muscle group. The peak of each curve indicates the point in the contraction cycle when each muscle group is computed to be located within the strongly contracted C bend. The ventral (V) group includes VO15, VO16, VA26 and VO28. The ventral lateral (VL) group includes VL12, DO5 and DL4 muscles. The dorsal lateral (DL) group includes DO11, DO19 and DO20. The dorsal group includes only muscle DL9. (L) indicates the left hemisegment, (R) denotes the right hemisgment. The horizontal dashed line denotes the approximate point in the contraction cycle where GCaMP6f signals are detectable in all 7 abdominal segments for the muscle group.

### A conceptual circuit for rolling emerges from the larval connectome

We next constructed a conceptual neuronal circuit for rolling that we inferred based on the muscles that we found to be involved in rolling. 22 motor neurons (MNs) innervate the 22 muscles that are most commonly involved in rolling on either the left or right side of the body. A recent study has mapped the synaptic connections of 236 premotor neurons (PMN) and 54 motor neurons (MN) insegment A1 of an abdominal ganglion larval connectome and 202 of these PMNs synapse directly onto MNs [2]. We found that 102 of these A1 PMNs synapse directly onto the 22 MNs that control the muscles involved in rolling. In order to conceptualize how these PMN-to-MN connections might give rise to a motor circuit for rolling behavior, we constructed a network graph representing the neurons and their connections (**Figure 4A**). Neurons in the network are represented as color-coded nodes where the colors indicate the location of the target muscle on the ventral-to-dorsal axis (cooler colors indicate ventral and warmer colors indicate dorsal). The connections between neurons are graphed as edges that are weighted by synaptic strength (number of synapses). Interestingly, the network graph reveals a circularity in the connectivity that roughly matches the circular sequence of muscle activation that travels around the circumference of the larva during rolling (**Figure 4A)**. In addition, the position of neurons within the circular network graph appeared to be correlated with their position along the ventral-to-dorsal axis. In contrast, when we graphed a network for the pool of MNs that were not involved in rolling, along with their PMN inputs, the circularity was not evident and MNs in the graph did not show a segregation according to their dorsal ventral location targets (**Figure S4)**.

**Figure 4:**
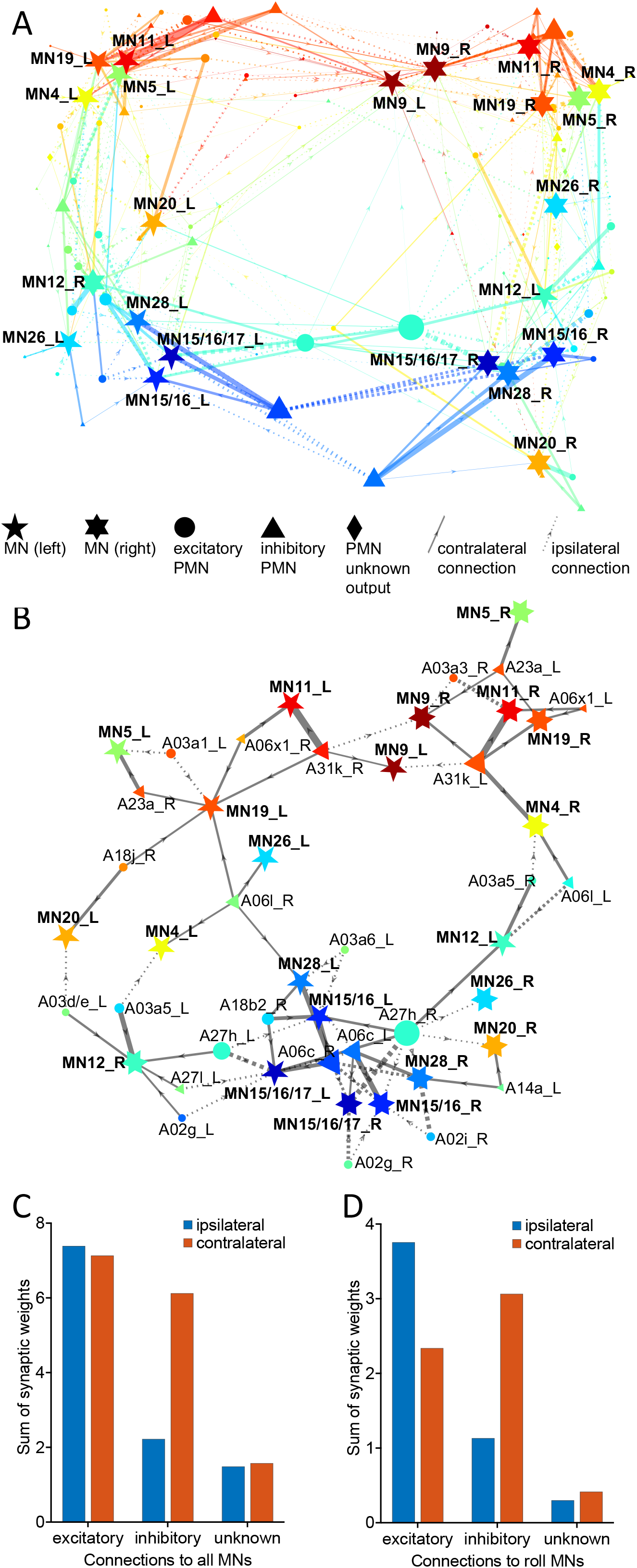
Putative connectivity network for PMNs and MNs involved in rolling. (A) Network graph representing synaptic connections from PMNs to MNs that target muscles that are activated during rolling. Node placement is determined automatically by a MATLAB algorithm based on the input matrix of connections between nodes. MNs are represented by pentagrams (left side) and hexagrams (right side). PMNs are represented by circles (excitatory synaptic output), triangles (inhibitory synaptic output), and diamonds (unknown synaptic output). Solid lines represent contralateral connections, dotted lines represent ipsilateral connections. The arrows represent the direction of the connection, which always goes from PMN to MN. The color of the MN nodes represents the location of their target muscle on the ventral-to-dorsal axis of the larva: blue for ventral, green for lateral, red for dorsal. The color of the PMN nodes and their output connections is an aggregate representation of their targets on the ventral-to-dorsal axis. The width of the lines represents the weight of the connections (determined by synapse number). The size of the PMN nodes represents the total weights of the outgoing connections. (B) Network graph representing the strongest synaptic connections of the network in panel A. A cutoff threshold for synaptic weights was chosen such that every MN would be connected to the network by at least one PMN connection. For visual clarity, PMNs that connect to only one MN were removed. Node placement was determined by a MATLAB force algorithm that spreads more distantly connected nodes farther apart. The shape, size, and color of the nodes and connecting lines are the same as in panel A. (C) The sum of synaptic weights between all PMNs in larval segment a1 and their target MNs, for excitatory, inhibitory, and unknown synapses to ipsilateral or contralateral connections. (D) The sum of synaptic weights between PMNs and MNs that target muscles that are activated during rolling, for excitatory, inhibitory, and unknown synapses to ipsilateral or contralateral connections.

In order to better visualize the stronger, and possibly most important, connections in the network, we set a threshold for the synaptic strengths, such that each MN would remain connected to the network by at least 1 edge. For visual clarity, we also removed any PMNs that connected to only a single MN. The circularity remains apparent in this simplified network graph and the circuit motifs of potentially important PMNs in this motor network can be more easily identified (**Figure 4B**).

We noted features of the graphed network that may be of importance for rolling locomotion. First, it appeared as if ipsilateral excitatory PMNs might be prominent. Second, it appeared as if contralateral inhibitory neurons were strongly connected to the network. These features are of interest because the muscle recruitment pattern for rolling that we characterized showed unilateral activation of each of the four coactive muscle groups in certain phases of the rolling cycle. This pattern could be facilitated by a combination of ipsilateral excitation and contralateral inhibition. Indeed, when we graphed the synaptic weights of the PMN inputs to the network we observed an excess of ipsilateral excitation relative to contralateral excitation (**Figure 4C**). We also found an excess of contralateral inhibition relative to ipsilateral inhibition (**Figure 4C)**. In contrast, PMN inputs to the motor network overall did not show the bias of ipsilateral excitation that was seen in the rolling motor network (**Figure 4D**). However, an excess of contralateral inhibition (relative to ipsilateral inhibition) is a general feature of the motor network that is found in both rolling and non-rolling (**Figure 4D**). Thus, we suggest that ipsilateral excitation may be important in driving the rolling network.

How and if the neurons within this network actually contribute to rolling behavior remains unknown. It is notable that several of the PMNs that are strongly connected in the network have been previously implicated in forward and/or reverse crawling. For instance, the inhibitory A31K neuron and the excitatory A27h neuron of the putative rolling network have both been shown to contribute to the crawling circuits [2,13]. If these neurons are also involved in rolling, then this would indicate polymodal function for neurons that are shared in the circuitry for rolling and crawling in the abdominal ganglion. Another question for the future is how the rolling motor network connects to the command like Goro neurons and other upstream pathways that trigger rolling [7–11]. Along these lines, it is interesting to note that the Goro command neurons have been reported to connect to the A10f interneuron which provides excitatory inputs into the A03a1 PMN that appears within our proposed rolling circuit [12]. While the vast majority of A10f outputs are unknown, three other PMNs in our putative rolling network (A06l, A07f2, A08e1) also receive its inputs [1]. Thus, the hypothetical network that we have proposed here provides a starting point that will enable further investigations on how the larval nervous system carries out a variety of distinct motor programs, such as crawling and rolling, with a single set of neurons.

## Acknowledgements

Important contributions in early pilot studies were made by Dr. Andrew Bellemer and Dr. Melanie Chin of the Tracey lab. We thank Dr. Aref Zarin and Dr. Chris Q. Doe for sharing of the R44H10-LexA muscle driver strain prior to publication. We thank members of the Tracey lab for helpful comments on the manuscript. We thank the Bloomington Drosophila Stock Center for fly strains. Funding to the study was provided by the National Institute of General Medical Sciences (WDT NIGMS 5R01GM086458), the Indiana University Foundation and the Linda and Jack Gill Center for Biomolecular Sciences.

## Supplementary Videos

Video S1. Recording of larval clockwise rolling. Related to Figure 2. Pseudocolor ranging from blue to red denote the GCaMP6f signals from low to high. Video was recorded at 60fps, the playback rate set as 10fps.

Video S2. Recording of larval counterclockwise rolling. Related to Figure 3. Pseudocolor ranging from blue to red denote the GCaMP6f signals from low to high. Video was recorded at 60fps, the playback rate set as 10fps in this video.

Video S3. Video of rotating 3D plot of Figure 2C. Related to Figure2.

Video S4. Video of rotating 3D plot of Figure 3C. Related to Figure3.

## STAR Methods

### Drosophila maintenance

Flies were maintained on standard Bloomington stock center cornmeal-agar food at 22°C in a relative humidity of 70%. A list of all the strains used in this study is detailed in the Key resources table.

### Rolling behavior imaging

Optogenetic activation with channelrhodopsin2-YFP in *Drosophila* larvae requires dietary supplementation with *all-transretinal* (ATR). We found that rolling of early second instar larvae could be facilitated by prefeeding the ATR to their virgin mothers (with yeast paste containing 1mM ATR) for 3-4 days (presumably by maternal loading of ATR into eggs). We crossed the ATR fed female R44H10-LexA LexAop-GCaMP6f/CyO flies to and male *ppk-GAL4 UAS-Chr2:YFP/CyO RFP*. After 3 days mating in vials containing yeast that was supplemented with ATR the flies were then transferred to the apple juice plate cages with ATR yeast paste and allowed to lay eggs overnight in the dark in a 25°C incubator (70% humidity). 2nd instar larvae were raised on the apple juice and ATR yeast paste until second instar (as scored by size of larvae).

3% agarose pads were cut into 10mm(L)X10mm(W) squares. A rounded hole in the center of the agarose pad was scooped out using a pipette tip. A small droplet of water was then used to fill the hole in the agar pad and a second instar larva of genotype R44H10-LexA LexAop-GCaMP6f/*ppk-GAL4 UAS-Chr2:YFP* was placed in the water filled depression, and then covered by 1.5mm coverslip. The preparation was then placed coverslip down in an inverted microscope (Zeiss LSM V Live). Imaging was conducted with excitation by a 100 mW 488 nm laser at 50-80% power. This laser excitation was effective at triggering rolling (in a small fraction of tested larvae) due to expression of Chr*2:YFP* in nociceptors. Time series images were acquired with a 10X/0.3 objective, a wide-open pinhole, and a long pass 505 emission filter at 60 frames per second for approximately 10 seconds. The thickness of an optical section in these imaging conditions was 119.5 microns and the larval diameter at this stage of development is approximately 250 microns. Our imaging conditions therefore allowed us to observe muscle activity in half the circumference of the larva at any given moment.

### The abdominal body wall muscle activity assay

The muscle activity was determined by the visual inspection of GCaMP6f signals from the time series recordings. Muscles were scored as ON based on a clearly observable rise in GCaMP6f fluorescence and clearly visible shortening of the visible muscle. The 8-bit grayscale images were processed with a rainbow pseudocolor scale in which colors from blue to red indicate intensity of GCaMP6f fluorescence. Inactive muscles were not scored as ON if their intensity resembled baseline of GCaMP6f fluorescence. Our imaging setting allows us to analyze individual muscle activity in half of the larvae in which muscles are visible and nearest to the coverslip.

### Data Analysis

The duration of a rolling cycle (ΔTime) was calculated as the time interval between two recording time points (at the start and end of a roll) when the larval ventral midline in A4 segment was centered in the image plane. The timestamp at the beginning of the rolling cycle (when segment A4 was centered on the ventral midline) was defined as TimeV. This allowed us to transform the true recording time to normalized time (NTime) from 0 to 1 as follows: NTime=(TimeR-TimeV)/ΔTime, where TimeR is a single timestamp from a frame in the recording and TimeV is defined above. At NTime=0 the ventral midline is centered, at NTime=0.5 the dorsal midline of the larva is centered, at NTime=1 the ventral midline is again centered after the completion of a 360 degree turn of a roll (**Figure S1**).

Our schematized estimate of the contraction cycle was made by first measuring the normalized position of each muscle relative to the ventral midline of the larva. An average position was then determined for each muscle group. We empirically determined the time point of the recordings when the muscle groups were located at the midline of the larva. The ventral (V) group includes VO15, VO16, VA26 and VO28 muscles. The ventral lateral (VL) group includes VL12, DO5 and DL4. The dorsal lateral (DL) group includes DO11, DO19 and DO20. The dorsal group includes only DL9. We defined the start point of contraction for the muscle group as the average for the normalized time point when the GCaMP6f signal began to rise in at least one of the abdominal segments in the muscles of a group. We assumed that the maximum contraction of the muscle group occurs when the muscle group was located within the C bend of the roll and that this position is a quarter cycle from the point when the muscle group is located at the midline. The estimation of the contraction cycle assumes a symmetrical pattern of contraction that continues beyond the point when all seven segments show GCaMP signals rising (dashed horizontal line in Figures 2E, 3E). Relaxation for each muscle group begins when the muscle group resides in the C bend (the peak of the curves) according to our calculations.

### Connectivity Network Methods

The directional graph networks were generated using the digraph object in MATLAB based on the connectivity matrix between PMNs and MNs published by Zarin et al. (2019). The connectivity matrix provides directional weighted connections from PMNs to MNs, where the connection weights are a percentage of the total synaptic inputs to each MN. Connections from PMNs to PMNs were not included. PMN nodes in the network graphs were limited to PMNs whose soma is located in segment a1. A network graph was generated for all segment a1 PMNs connected to a1 MNs that target muscles activated during rolling (Figure 4A), and for all segment a1 PMNs connected to a1 MNs that target muscles not involved in rolling (Figure S2A). The graphs were plotted using the MATLAB plot function, and placement of the nodes was automatically determined by MATLAB using the ‘subspace’ layout. The color of the MN nodes represents the location of their target muscles on the ventral-to-dorsal axis of the larva. The colors were selected at equally spaced intervals from the jet colormap in MATLAB from blue (ventral) to red (dorsal). The color of the PMN nodes represents the combination of their MN targets, such that PMNs with mostly ventral connections are an intermediate shade of blue, while PMNs with mostly dorsal connections are a shade of red, and PMNs with lateral or broadly mixed connections are closer to green or orange. The network graphs were simplified to show only the strongest connections (Figures 4B and S2B). Cutoffs for the connection weights were thresholded to show the minimal number of connections while still keeping each MN connected to the network by at least one PMN connection. The cutoff for the simplified network to muscles activated during rolling was 0.0385, the cutoff for the simplified network to muscles not involved in rolling was 0.0225. For visual clarity, PMNs connected to only one target MN were also removed. The simplified networks were plotted in MATLAB using the ‘force’ layout which spaces distantly connected nodes further apart and increases visual clarity.

## Key Resources Table

| Reagent or Resource | Source | Identifier |
| --- | --- | --- |
| Experimental Models:<br>Organisms/Strains |  |  |
| *Drosophila*: *R44H10-LexA LexAop-GCaMP6f/CyO* | Zarin et al | N/A |
| *Drosophila*: *ppk-GAL4 UAS-ChR2/CyO RFP(mCherry)* | This study | N/A |
| Software and algorithms |  |  |
| ImageJ | Imagej.net/contributors | ImageJ1.52a |
| MATLAB | MathWorks | R2021b |
| Adobe Illustrator | Adobe | CC2022 |
| Excel | Microsoft | 16.0.4738.1000 |
| Zen 2009 | Zeiss | 6,0,0,303 |

## Notes

### Competing Interest Statement

The authors have declared no competing interest.

