## Supplemental Figures for "The motor pattern of rolling escape locomotion in *Drosophila* larvae"

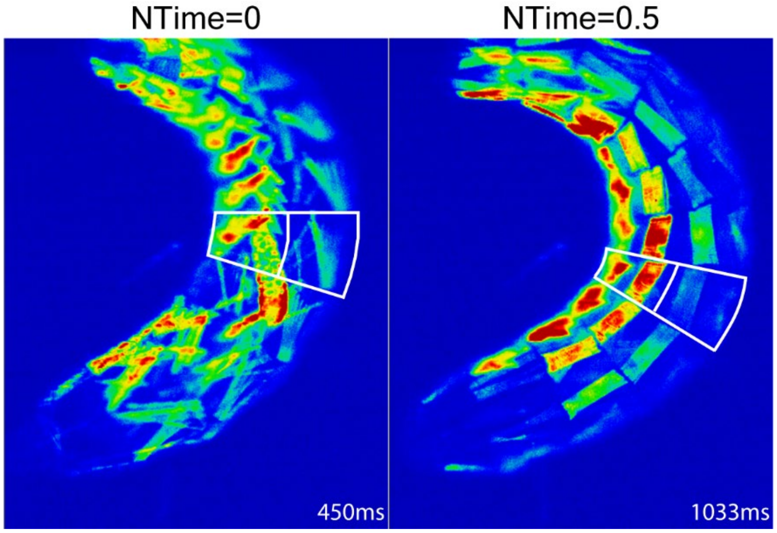


Figure S1: Larval rolling phase

NTime=(TimeR-TimeV)/ΔTime, where NTime is normalized time and TimeR is a single timestamp from a frame in the recording. The duration of a rolling cycle (ΔTime) was calculated as the time interval between two recording time points (at the start and end of a roll) when the larval ventral midline in A4 segment was centered in the image plane. The timestamp at the beginning of the rolling cycle (when segment A4 was centered on the ventral midline) was defined as TimeV. At NTime=0 the ventral midline is centered, NTime=0.5, the dorsal midline of the larva is centered.

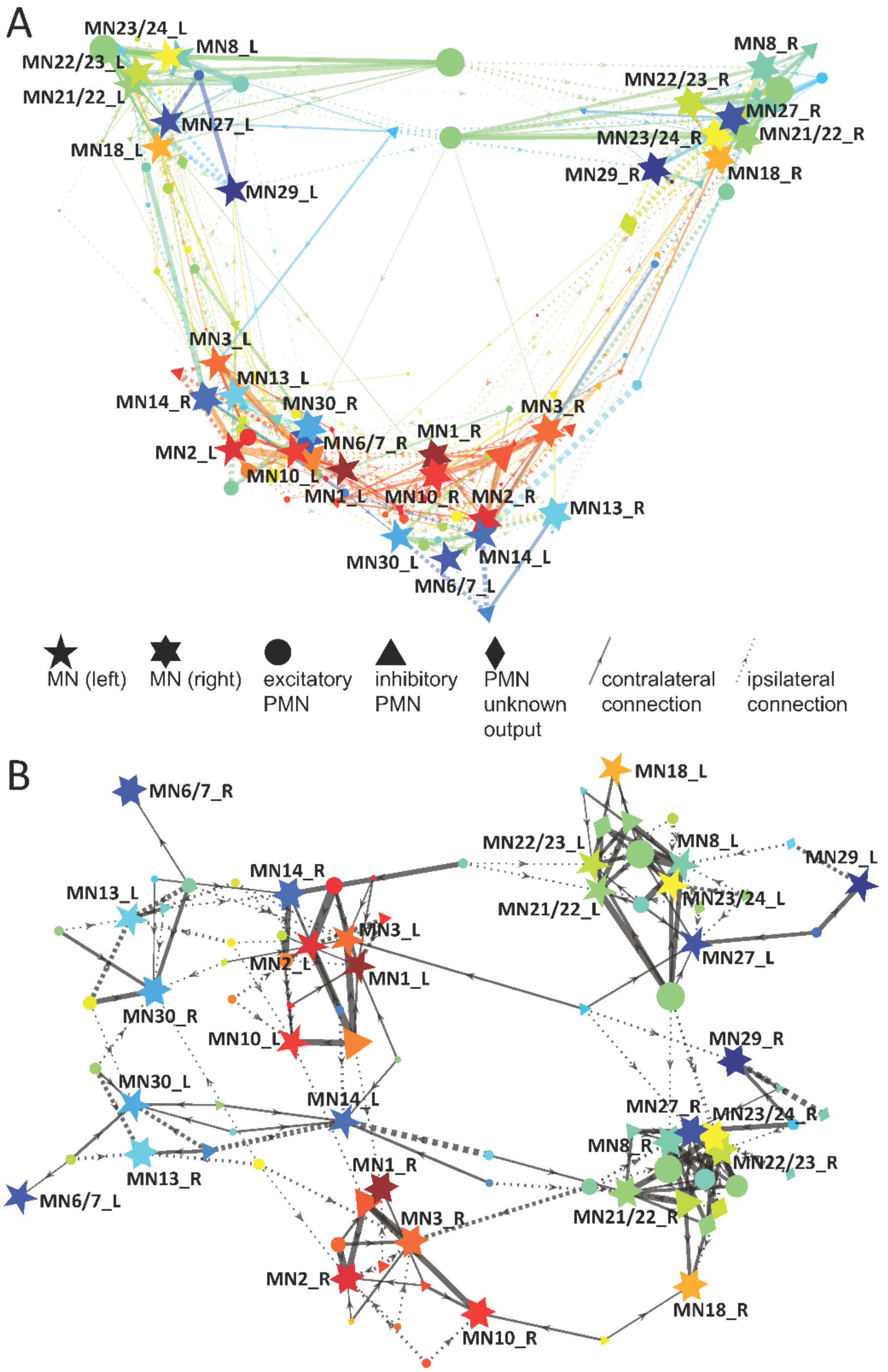


Figure S2: Conceptual connectivity network for PMNs and MNs that target muscles that are not involved in rolling.

(A) Network graph representing synaptic connections from PMNs to MNs that target muscles that are not activated during rolling. Node placement is determined automatically by a MATLAB algorithm based on the connectivity matrix between nodes. MNs are represented by pentagrams (left side) and hexagrams (right side). PMNs are represented by circles (excitatory synaptic output), triangles (inhibitory synaptic output), and diamonds (unknown synaptic output). Solid lines represent contralateral connections, dotted lines represent ipsilateral connections. The arrows represent the direction of the connection, which always goes from PMN to MN. The color of the MN nodes represents the location of their target muscle on the ventral-to-dorsal axis of the larva: blue for ventral, green for lateral, red for dorsal. The color of the PMN nodes and their output connections is an aggregate representation of their targets on the ventral-to-dorsal axis. The width of the lines represents the weight of the connections (determined by synapse number). The size of the PMN nodes represents the total weights of the outgoing connections.

(B) Network graph representing the strongest synaptic connections of the network in panel A. A cutoff for synaptic weights was chosen such that every MN would be connected to the network by at least one PMN connection. For visual clarity, PMNs that connect to only one MN were also removed. Node placement is determined by a MATLAB force algorithm that spreads distantly connected nodes apart. The shape, size, and color of the nodes and connecting lines are the same as in panel A.
